# Chemical inhibition of PIN auxin transporters by the anti-inflammatory drug Naproxen

**DOI:** 10.1101/2022.10.13.512040

**Authors:** Jing Xia, Mengjuan Kong, Zhisen Yang, Lianghanxiao Sun, Yakun Peng, Hong Wei, Wei Ying, Yongxiang Gao, Jiří Friml, Xin Liu, Linfeng Sun, Shutang Tan

## Abstract

The phytohormone auxin plays central roles in many growth and developmental processes in plants. Development of chemical tools targeting the auxin pathway is useful for both plant biology and agriculture. Here we uncover that Naproxen, a synthetic compound with anti-inflammatory activity in humans, acts as an auxin transport inhibitor targeting PIN transporters in plants. Physiological experiments indicate that exogenous Naproxen treatment affects pleiotropic auxin-regulated developmental processes. Further cellular and biochemical evidence supports that Naproxen suppresses auxin transport, specifically PIN-mediated auxin efflux. Moreover, biochemical and structural analysis confirms that Naproxen binds directly to PIN1 protein, via the same binding cavity as the IAA substrate. Thus, by combining cellular, biochemical, and structural approaches, this study well establishes that Naproxen is a PIN inhibitor and elucidates the underlying mechanisms. Further use of the compound may advance our understanding on the molecular mechanisms of PIN-mediated auxin transport, and expand our toolkit in auxin biology and agriculture.

## Introduction

Plant growth and development requires coordination of internal signals and environmental cues. The phytohormone auxin (indole-3-acetic acid, IAA) plays fundamental roles in almost every aspect of plants’ life (Friml, 2022). Recent decades of genetic studies have well established the molecular framework for auxin biosynthesis (Zhao, 2018), transport (Adamowski and Friml, 2015; Tan et al., 2021), and signaling (Lavy and Estelle, 2016; Gallei et al., 2020), cooperating together for multitude of auxin functions.

Specifically, IAA is synthesized by the TAA1/TAR and YUCCA (YUC) enzymes from the tryptophan precursor, and additionally, multiple modifications control IAA homeostasis in plants (Zhao, 2018). The nuclear and cytosolic TRANSPORT INHIBITOR RESPONSE1 (TIR1)/AUXIN SIGNALLING F-BOX (AFB)-AUXIN/INDOLE-3-ACETIC ACID (AUX/IAA) pathway mediates auxin function in transcriptional regulation (Salehin et al., 2015; Lavy and Estelle, 2016) and also in non-transcriptional rapid responses related to root growth regulation (Li et al., 2022). On the other hand, the cell-surface AUXIN BINDING PROTEIN1 (ABP1)-TRANSMEMBRANE KINASEs (TMKs)-based auxin perception confers a rapid cellular auxin responses through global modulation of the phosphoproteome (Friml et al., 2022). Notably, the directional auxin transport between cells, namely polar auxin transport (PAT), is the key feature of auxin action. Plasma membrane (PM)-localized PIN-FORMED (PIN) family auxin transporters pump auxin out of the cell, playing essential roles in PAT (Petrášek et al., 2006). Intriguingly, PIN transporters reside unequally at the PM, and the polarity of PIN proteins determines the directionality of intercellular auxin flow, which is crucial in many patterning processes as well as in the asymmetric growth during tropic responses (Adamowski and Friml, 2015; Tan et al., 2021; Konstantinova et al., 2022). Moreover, AUXIN1 (AUX1)/LIKE AUXINs (LAXs) are auxin importers (Marchant et al., 1999), and ABC transporters have been reported to be also involved in auxin export (Serrano et al., 2013; Hao et al., 2020). Given the central roles of auxin in plant growth and development, exogenous auxin applications and genetic or pharmacological manipulations of the auxin action are widely used in both plant research and agriculture.

In the history of auxin research, various chemical tools have been developed and they play important roles in overcoming the problem of functional redundancy for gene families (Kong et al., 2022). For example, L-Kynurenine targets TRYPTOPHAN AMINOTRANSFERASE OF ARABIDOPSIS (TAA1)/ TRYPTOPHAN AMINOTRANSFERASE RELATED PROTEIN (TAR) enzymes to inhibit auxin biosynthesis *in planta* (He et al., 2011). Auxinole is an auxin antagonist of TIR1/AFB-AUX/IAA mediated auxin signaling. As for auxin transport, there are a family of compounds namely auxin transport inhibitors (ATIs), of which, N-1-Naphthylphthalamic acid (NPA) is widely used. Application of NPA to plants leads to various defects (e.g., interference of phototropism and gravitropism, inhibition of lateral root formation, suppression of vascular development, and deficiency in shoot development) due to suppression of auxin transport across cells. Recent biochemical and structural studies reveal that NPA directly binds to and inhibits the activity of PIN-FORMED (PIN) auxin transporters (Abas et al., 2021; Teale et al., 2021; Lam Ung et al., 2022; Su et al., 2022; Yang et al., 2022). Interestingly, NPA binds to the same pocket as the PIN substrate IAA, suggesting that NPA is a competitive inhibitor of PIN proteins, which is a paradigm shift in our understanding of NPA action. Thus, identification of novel PIN inhibitors will be useful in both plant biology and agriculture.

Non-steroidal anti-inflammatory drugs (NSAIDs), such as aspirin and ibuprofen, are a set of compounds which are used to relief pain and fever in humans (Huls et al., 2003; Duggan et al., 2011). Among NSAIDs, salicylic acid (SA) is an ancient drug, and its use in reducing pain and fever can be tracked to the Hippocrates era. Indeed, SA is an immune signal perceived by the NON-TRANSCRIPTION OF PR GENES (NPR) family receptors to regulate pathogen responses in plants (Fu et al., 2012; Wu et al., 2012; Ding et al., 2018; Wang et al., 2020). Recent studies show that SA has also been found to regulate plant growth and development (Du et al., 2013; Rong et al., 2016; Pasternak et al., 2019; Ke et al., 2021) via modulating clathrin-mediated endocytosis (Du et al., 2013) and/or targeting the A subunits of Protein Phosphatase 2A (PP2A) (Tan et al., 2020b). Oxicam-type of NSAIDs were reported to target NPR1 proteins-mediated immune pathway as antagonists of SA in plants (Ishihama et al., 2021). Our previous studies showed that synthetic non-steroidal anti-inflammatory drugs (NSAIDs) showed activity in plant cells (Tan et al., 2020c). Notably, amongst other NSAIDs, meclofenamic acid (Meclo) and flufenamic acid (Fluf) target the immunophilin-like protein TWISTED DWARF1 (TWD1, also called FKBP42) to regulate the dynamics of actin cytoskeleton and subsequent endomembrane trafficking (Tan et al., 2020c). Nonetheless, not all NSAIDs were detected to bind the FKBP domain of TWD1 from surface proton resonance (SPR) assays and hence how the other NSAIDs are involved in regulation of plant growth and development is unclear.

In this report, we describe that Naproxen [(S)-(+)-6-Methoxy-α-methyl-2-naphthaleneacetic acid, Extended Data Fig. 1a], a common NSAID, exhibits a typical auxin transport inhibitor activity. Biochemical and structural experiments reveal that Naproxen binds directly to PIN1 auxin transporter and inhibits its activity. Therefore, Naproxen is a novel auxin transport inhibitor, with potential to be used in both auxin research and agriculture.

## Results

### Naproxen treatment modulates *Arabidopsis* growth and development

NSAIDs, represented by aspirin and ibuprofen, are widely used medicines to treat various aches and inflammations (Huls et al., 2003; Yiannakopoulou, 2013). In our previous study, it was shown that NSAIDs share bioactivities in plant cells and that, amongst other NSAIDs, Meclo and Fluf target TWD1 immunophilin-like protein to modulate the dynamics of actin filament and endomembrane trafficking, thus regulating trafficking of PIN2 protein to shape root growth and development in *Arabidopsis*. To further verify the molecular mechanisms underlying other NSAID chemicals functioning in plants, herein we focused on one of them, Naproxen, and examined its physiological effects in detail. Naproxen is widely used as an anti-inflammatory drug, to relief pain and swelling by inhibiting the Cyclooxygenase-2 (COX-2) enzyme to reduce prostaglandin production in animal cells (Duggan et al., 2010). When growing on Murashige & Skoog (MS) plates supplemented with Naproxen (10, 20, 40 µM, respectively), *Arabidopsis* seedlings exhibit shorter (Fig. 1a, b) and less gravitropic primary roots (Fig. 1a, d, and Extended Data Fig.1c), as well as reduced number of lateral roots as compared with those on the DMSO control (Fig. 1c, and Extended Data Fig.1b). In summary, these effects of Naproxen on root development are auxin-related, reminiscent of that of NPA, a widely used auxin transport inhibitor, which was used as a positive control. Thus, we explored the activities of Naproxen in auxin transport-dependent biological processes in more detail.

**Figure 1.**
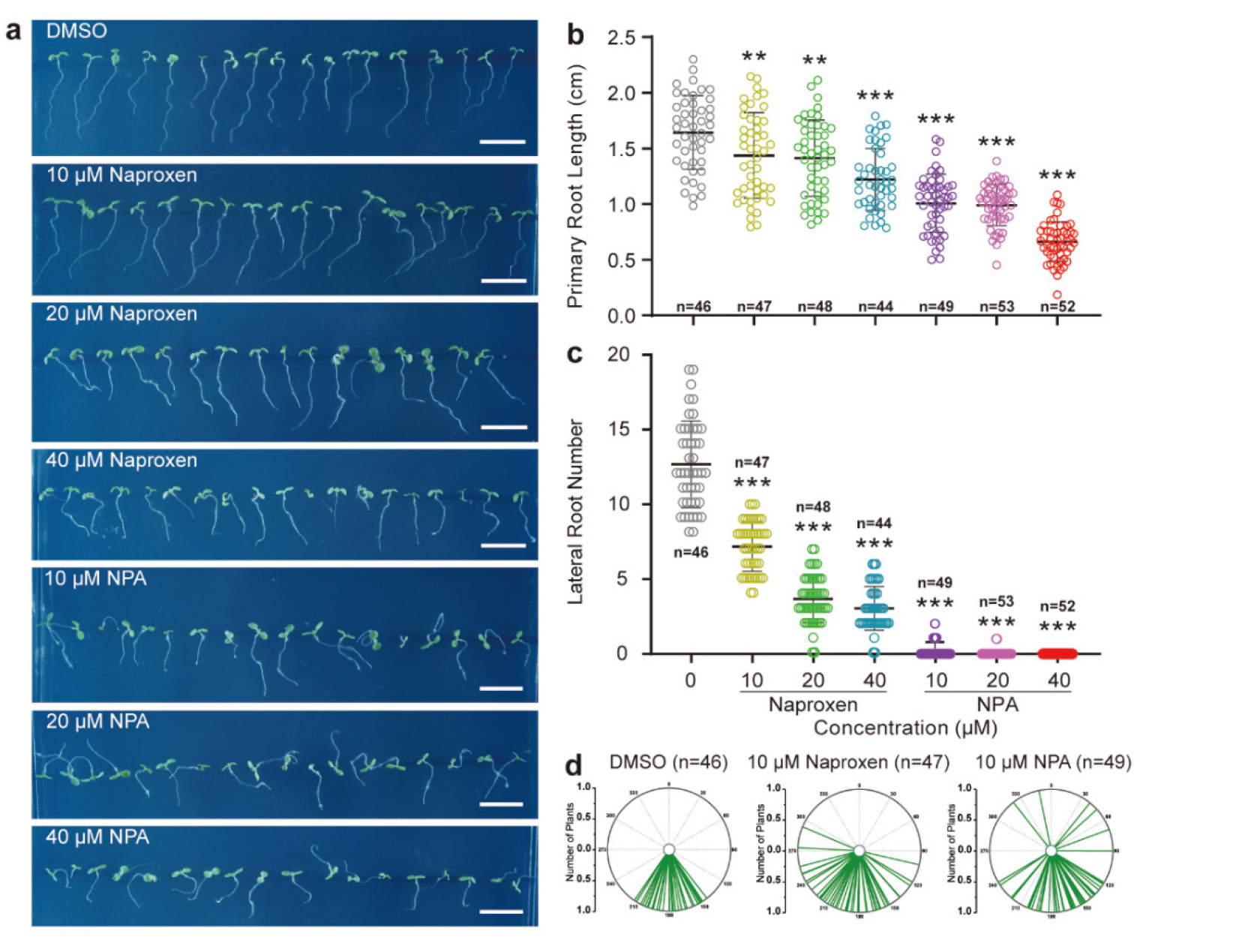
Physiological effects of Naproxen on *Arabidopsis* root development. **a**, Representative images showing the morphological changes of seven-day Col-0 seedlings grown on MS medium supplemented with Naproxen and NPA at indicated concentrations. DMSO as the solvent control. Scale bars, 1 cm. **b**, Dose-dependent effect of Naproxen and NPA inhibiting primary root elongation. Seven-day-old Col-0 seedlings. DMSO as the solvent control. **c**, Naproxen and NPA suppressed lateral root formation. The emerged lateral roots of ten-day-old Col-0 seedlings treated with Naproxen and NPA were counted; n = 46, 47, 48, 44, 49, 53, and 52 seedlings, respectively. **d**, Naproxen and NPA interfered with root gravitropism. Seven-day-old Col-0 seedlings grown constantly on MS medium supplemented with DMSO, Naproxen and NPA, as indicated. Each line represents the root tip angle of 1 individual seedling in polar bar charts. *p* values were calculated by comparing different treatments to DMSO with an unpaired *t* test with Welch’s correction, \*\**P*<0.01, \*\*\**P*<0.001.

Auxin plays a central role in plant tropic responses toward gravity and light, namely gravitropism and phototropism respectively. For both processes, PIN-mediated polar auxin transport determines the asymmetric (re)distribution of this growth hormone, thus ensuring differential growth of plant organs (Friml et al., 2002; Ding et al., 2011; Rakusová et al., 2016). When wild-type Col-0 *Arabidopsis* seedlings were grown on MS medium with the DMSO solvent control under dark, they grew upright with a long hypocotyl and a closed apical hook (so-called skotomorphogenesis), whereas seedlings on MS plates with Naproxen are less gravitropic (Extended Data Fig. 2a, b). Upon gravistimulation by turning 90 degrees, seedlings bent up on media with the DMSO control, while those on MS medium with Naproxen failed, suggesting a defect in either gravity perception or gravity-stimulated differential growth (Extended Data Fig. 2c, d). Moreover, etiolated seedlings failed to form a normal apical hook, which is strictly dependent on auxin activity (Zadnikova et al., 2010; Cao et al., 2019), on MS plates supplemented with Naproxen or NPA (Extended Data Fig. 2). Similarly, Naproxen also compromised phototropism of *Arabidopsis* seedlings (Extended Data Fig. 3a, b). These results suggest that Naproxen impaired numerous processes of auxin-mediated plant growth and development, in a similar way as the traditional auxin transport inhibitor NPA.

### Naproxen interferes with auxin response *in planta*

To test if Naproxen interferes with the auxin pathway, we employed the *DR5rev::GFP* auxin responsive reporter (Friml et al., 2003). Auxin forms a regular maxima at the root tip, which is important for root meristem patterning and gravitropic response (Adamowski and Friml, 2015). Decreased activity of PIN transporters in the responsible activating kinase mutants such as *d6pk d6pkl1 d6pkl2* and *pdk1*.*1 pdk1*.*2* caused an expansion of root tips (Tan et al., 2020a; Xiao and Offringa, 2020). Similar phenomenon is also observed in NPA-treated roots, with a broader DR5 signal (Fig. 2a). Intriguingly, Naproxen exhibits the same effect though a relatively higher concentration (40 µM) than that of NPA (10 µM) (Fig. 2a). This constitutive treatment also led to tissue swelling in root tips with overgrowth of gravity-sensing columella cells (statocytes) revealed by Lugol staining (Fig. 2b). The over-proliferation of statocytes was correlated to the DR5 pattern, which was reported to be due to exaggeration of the region with auxin maxima (Nakamura et al., 2019). This typical effect of NPA prompted us to test if Naproxen functions in a similar way.

**Figure 2.**
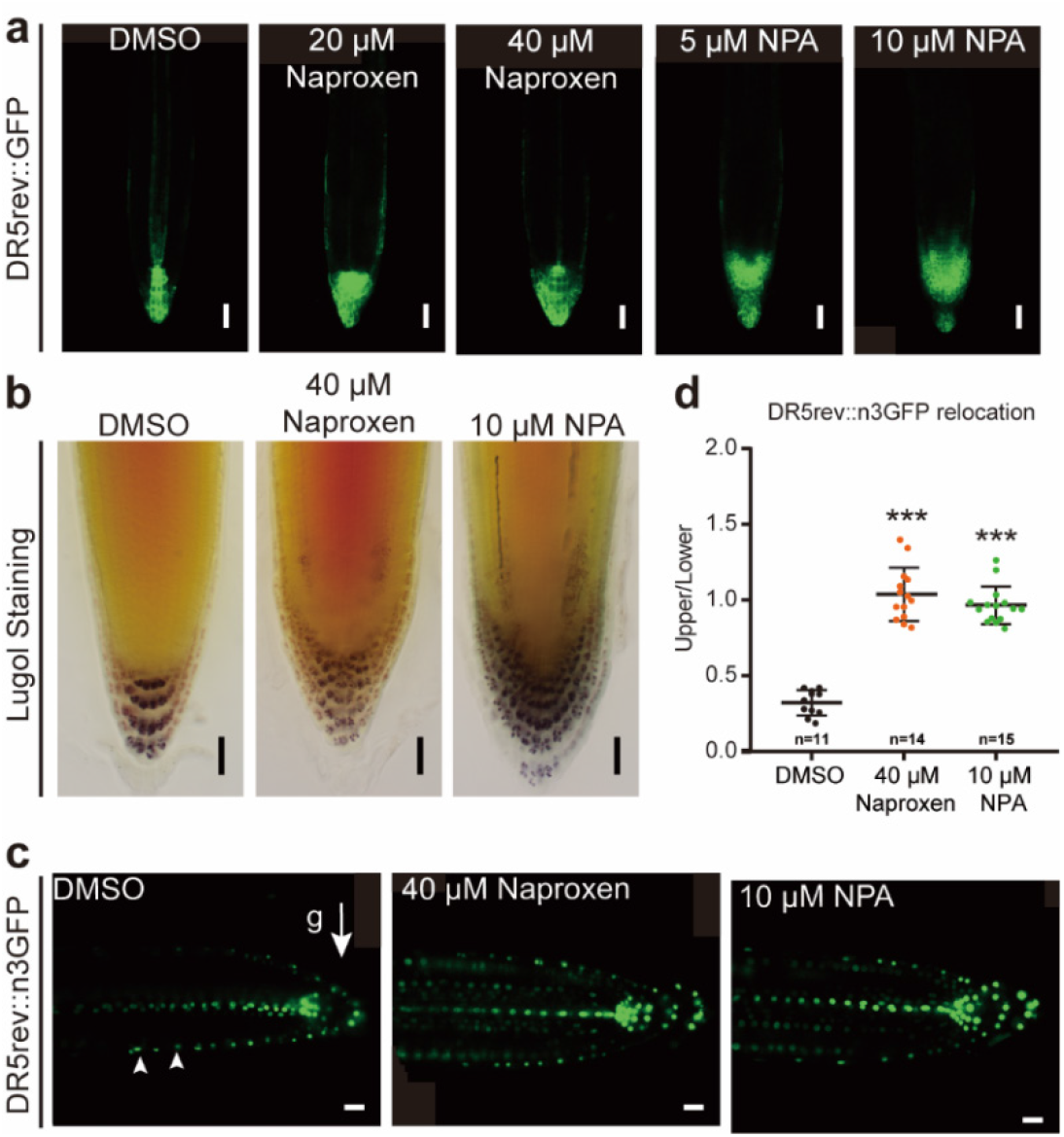
Naproxen interferes with auxin response *in planta*. **a**, The regular auxin-responsive pattern of *DR5rev::GFP* was disrupted by Naproxen and NPA treatments. Five-day-old *DR5rev::GFP* seedlings grown on plates containing Naproxen and NPA with indicated concentrations were imaged with confocal laser scanning microscopy (CLSM), n = 12-16, and scale bars, 50 μm. **b**, Constant Naproxen and NPA treatments changed the pattern of statocytes. Five-day-old Col-0 seedlings under indicated treatments were stained with Lugol’s solution and then imaged with a differential interference contrast (DIC) microscope, n = 15-20, and scale bars, 20 μm. **c** and **d**, Naproxen and NPA treatments suppressed the re-distribution of DR5rev::n3GFP pattern under gravi-stimulation. Five-day-old *DR5v2::ntdTomato;DR5rev::n3GFP* seedlings grown on normal plates were transferred to DMSO/Naproxen/NPA-containing plates, and turned by 90°. Root tips were imaged with CLSM for the GFP channel. **c**, Representative images. The arrow head indicates the direction of gravity. Scale bars, 20 μm. **d**, The upper:lower ratio for the DR5 signal was measured to indicate the relocation; n = 11, 14, 15, respectively. p values were calculated by comparing different treatments to the DMSO control with an unpaired *t* test with Welch’s correction, \*\*\**P*<0.001.

In the gravitropic response, auxin (as monitored by DR5) re-distributes to accumulate at the lower side of the root, where it inhibits cell elongation to promote the root to bend down (Baster et al., 2013). As expected, Naproxen treatment impaired seedlings’ ability to respond to gravistimulation, along with a defect in asymmetric DR5 signal formation (Fig. 2c and d), suggesting that Naproxen might interfere with polar transport auxin from root tips during gravitropic response. Taken together, these results indicate that Naproxen might inhibit auxin transport in plant cells.

### Naproxen inhibits auxin transport via binding directly to PIN proteins

These above physiological and cellular analyses suggest that Naproxen might be an auxin transport inhibitor. With advantage of an auxin efflux assay based on HEK293F cells and purified PIN1 protein (Yang et al., 2022), we tested if Naproxen targets PIN auxin transporters directly. In the auxin efflux assay, which measured residual ^3^H-labelled radioactive IAA ([^3^H]IAA) in cells after transfer into isotope-free buffer, we found that Naproxen treatment led to a higher retention level of [^3^H]IAA in cells expressing PIN1 protein plus the activating kinase D6PK than the DMSO control, suggesting an inhibition of auxin export (Fig. 3a). This result implies that Naproxen may target PIN1 protein directly.

**Figure 3.**
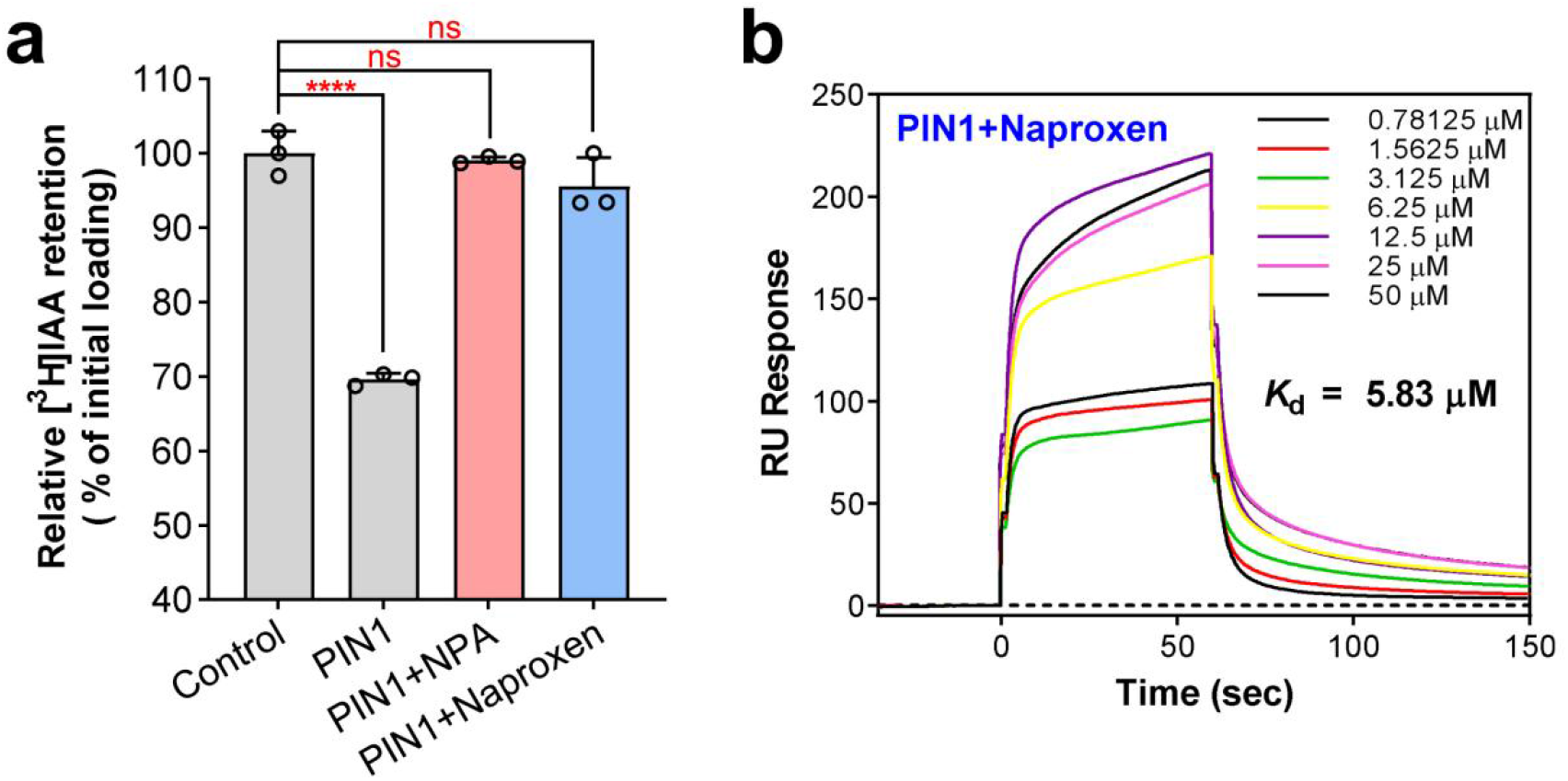
Naproxen binds to and inhibits the auxin efflux activity of PIN1. a, [^3^H]IAA retention relative to the initial loading in the absence or presence of NPA and Naproxen. Each data point is the average of three independent experiments. Statistical analysis was performed using one-way ANOVA with Dunnett’s multiple comparisons test. ns = not significant, **** *P* < 0.0001 for wild type versus control. Data are presented as mean ± s.e.m. b, The binding affinity between PIN1 and Naproxen as measured by SPR. RU, resonance units. The dissociation constant *K*_d_ is 5.83 μM.

To further test this hypothesis, we measured the binding between PIN1 and Naproxen using the surface plasmon resonance (SPR) assay, and results indicated that Naproxen indeed bound to PIN1 protein, with a kinetic dissociation constant (*K*_d_) value of approximately 5.83 µM (Fig. 3b). It is noteworthy that Naproxen has lower binding affinity to PIN1 protein than that of NPA, whose *K*_d_ value is 0.0166 µM as also revealed by SPR (Extended Data Fig. 4a). This corelates well with the physiological and cellular experiments that Naproxen exhibits a relatively lower activity than NPA. Furthermore, we also tested the binding affinity between Naproxen and other PIN members, PIN2, PIN5 and PIN8 specifically, respectively. The results show that Naproxen can bind to all PINs examined, with *K*_d_ values similar to that of Naproxen-PIN1 (Extended Data Fig. 4b-d).

### Structural basis for Naproxen inhibiting PIN-mediated auxin transport

To gain insight into the inhibition mechanism of Naproxen, we tried to determine the complex structure of PIN1 and Naproxen. Naproxen was added to the purified PIN1 protein bound with a synthetic nanobody, sybody-21 (Yang et al., 2022), prior to cryo-sample preparation. After cryo-EM data collection and processing, an EM map with an overall resolution of 3.7 Å was obtained (Fig. 4a, Extended Data Fig. 5b-g and Extended Data Fig. 6a). The overall EM density was similar to those of determined PIN1 structures. In the intracellular cavity, a clear density was observed which docked well with Naproxen (Fig. 4b and Extended Data Fig.6b). Such a cavity has also been revealed to accommodate the natural auxin IAA and the inhibitor NPA (Yang et al., 2022). In line with the result that Naproxen has a higher binding affinity to PIN1 protein (with a *K*_d_ value of 5.83 µM, Fig. 3b) than the substrate IAA (with a *K*_d_ value of 186 µM, as revealed by SPR in Yang et al., 2022), suggesting that Naproxen may be a competitive inhibitor of IAA for PIN transporters.

This complex structure aligns well with the NPA-bound structure of PIN1, with a root mean square deviation (r.m.s.d.) of 0.53 Å (672 Cα atoms aligned), both exhibiting an inward-facing conformation (Extended Data Fig.6). Like NPA, Naproxen binds to PIN1 through both hydrogen bonding and hydrophobic interactions (Fig. 4c, d). The carboxyl group of Naproxen lies between N112 and Y145 and forms a hydrogen bond with Y145. The naphthalene moiety is stacked between hydrophobic residues of PIN1, including V51, V115, I582 and V583 (Fig. 4c, d). This suggests a conserved binding mode between Naproxen or NPA and PIN1, which is also reflected in the well-superposed binding sites in the structures (Fig. 4e). However, the carboxyl group of NPA forms more hydrogen bonding with PIN1 like with the backbones of I582 and V583, and the benzene ring also forms hydrophobic stacking with Y145, L114 and V583 (Yang et al., 2022), which may contribute to the higher binding affinity with PIN1 than that of Naproxen.

**Figure 4.**
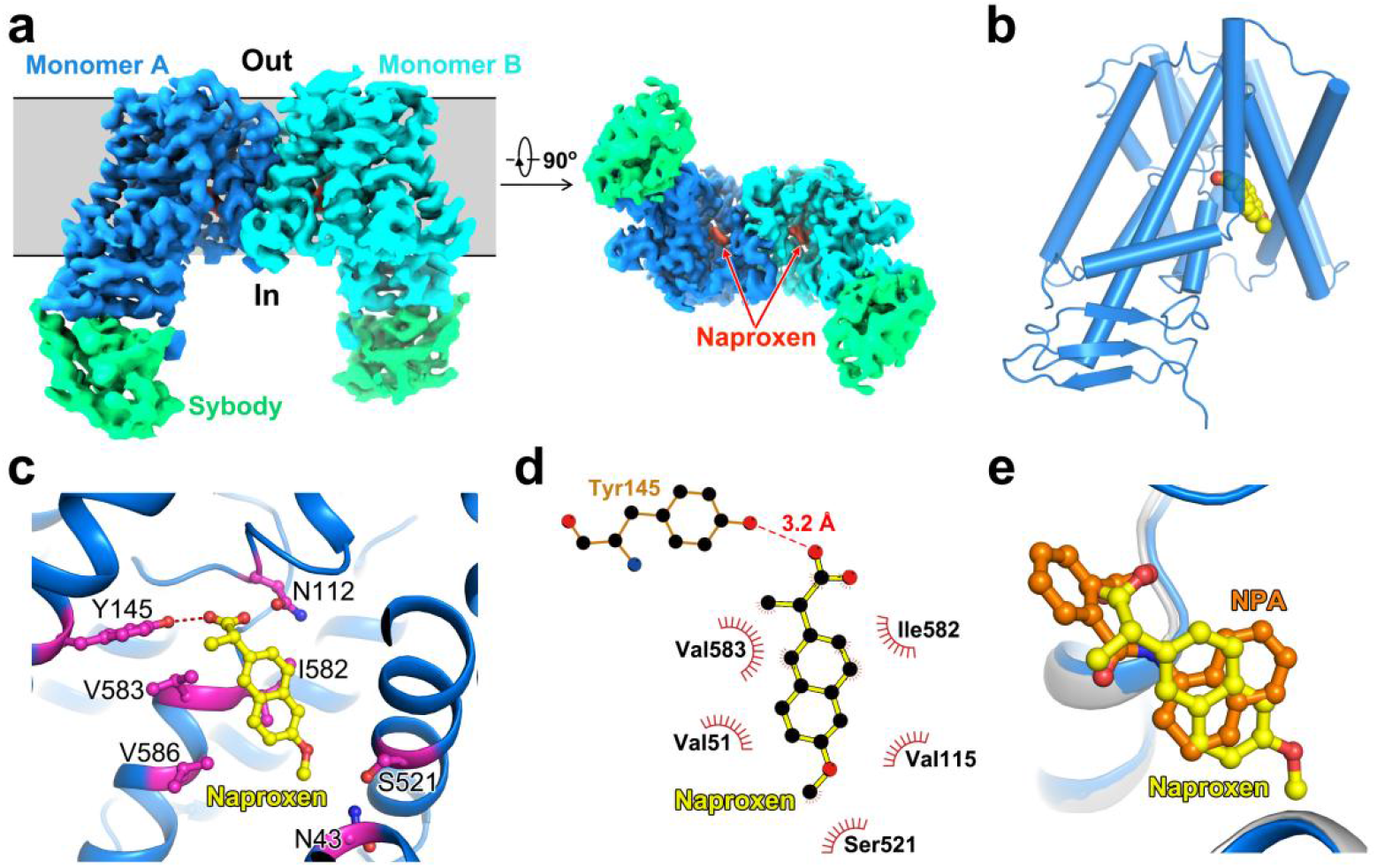
Structure of the Naproxen-bound PIN1. a, Overall cryo-EM map of PIN1 in complex with Naproxen. Densities of the two protomers are shown in blue (monomer A) and cyan (monomer B), respectively. The sybodies are colored green. Naproxen (red density) in the map of each PIN1 subunit are indicted with red arrows. Side view and bottom view are presented. b, Cartoon representation of PIN1 monomer bound Naproxen. Naproxen is shown as spheres. The structure of Naproxen-bound PIN1 exhibits an inward-facing conformation. c, Coordination of Naproxen by PIN1. Naproxen and interacting residues are shown in sticks. Hydrogen bonds are shown as red dashed lines. d, Schematic representation of the Naproxen-PIN1 interactions as shown by Ligplot^+^ (red dashed line, hydrogen bond; spokes, hydrophobic interactions). e, Superposition of the Naproxen-bound and NPA-bound (PDB code: 7Y9U) structures of PIN1. Naproxen and NPA are shown in sticks and colored yellow and orange, respectively.

## Discussion

Auxin plays essential roles in plant growth and development. Synthetic auxins and anti-auxin compounds are important tools in both plant research and modern agriculture. Amongst these chemicals, ATIs were used as herbicides in agriculture. In this study, we uncovered that Naproxen is a novel auxin transport inhibitor targeting directly PIN auxin transporters. Naproxen treatment leads to a variety of morphogenetic changes in *Arabidopsis* seedlings that can be ascribed to disturbance in auxin distribution. The effects of Naproxen in plants are also reminiscent of *pin* mutants, such as shortened root length in *pin1 pin3 pin4* or *pin1 pin3 pin4 pin7* (Friml et al., 2003), the lateral root deficiency in *pin1* (Benková et al., 2003), agravitropic roots in *pin2* (Chen et al., 1998; Luschnig et al., 1998) or *pin3* (Kleine-Vehn et al., 2010), agravitropic hypocotyls in *pin3 pin4 pin7* etiolated seedlings (Rakusová et al., 2016), and impaired phototropism in *pin3* or *pin3 pin4 pin7* mutants (Friml et al., 2002; Ding et al., 2011). Biochemical and structural analyses confirmed that Naproxen binds directly to the IAA-binding pocket of PIN1 protein. Intriguingly, although the binding profile is quite similar for NPA and Naproxen, there are still differences in the detailed interactions and binding affinities, suggesting potential for further engineering to develop novel compounds targeting PIN transporters.

We used purified PIN1 protein (Yang et al., 2022) to perform biochemical and structural analyses. However, given that Naproxen exhibits broader ATI activities *in planta* than PIN1 does, and that the active sites are conserved across PIN family auxin transporters, Naproxen may target other PIN homologues in the same manner, as supported by the SPR results (Extended Data Fig. 4b-d). Moreover, as we have noticed that other NSAIDs, such as indomethacin and ketoprofen, also preserve similar activities (i.e., causing enlarged root tips and expansion of DR5 signal) as Naproxen and NPA, it is likely that more NSAIDs also target PIN proteins (Tan et al., 2020). Notably, Naproxen is a widely used anti-inflammatory drug in humans, and it is safe for human health. This feature is crucial if Naproxen is further considered to be used as an herbicide in agriculture.

## Acknowledgements

We thank the Cryo-EM Center of the University of Science and Technology of China for the EM facility support. We are grateful to Y. Gao and all the other staff members for their technical support on cryo-EM data collection. This work was supported by the start-up fundings from University of Science and Technology of China (KY9100000026, KY9100000051, and KJ2070000079 to ST) and the Strategic Priority Research Program of Chinese Academy of Sciences (XDB37020103 to L.S.), National Natural Science Foundation of China (31900885 to X.L., 31870732 to L.S.), Natural Science Foundation of Anhui Province (2008085MC90 to X.L., 2008085J15 to L.S.), the Fundamental Research Funds for the Central Universities (WK9100000021 to S.T., and WK9100000031 to L.S.), and the USTC Research Funds of the Double First-Class Initiative (YD9100002016 to S.T., and YD9100002004 to L.S.). L.S. is supported by an Outstanding Young Scholar Award from the Qiu Shi Science and Technologies Foundation, and a Young Scholar Award from the Cyrus Tang Foundation.

## Author Contributions

X.L., L.S., and S.T. conceived the project and designed the experiments. The physiological effect of Naproxen was originally discovered by S.T. in J.F. lab at ISTA. M. K., Lianghanxiao S. and Y.P. performed physiological and cellular experiments under supervision of S.T.. Z.Y. and J.X. performed protein purification, biochemical assays, and structure determination work under supervision of X.L. and L.S.. Z.Y., J.X., H.W. and W.Y. performed the auxin transport assays. Z.Y. and Y.G. performed the cryo-EM data collection. All authors contributed to data analysis. J.F. contributes to manuscript preparation. X.L., L.S., and S.T. wrote the manuscript with inputs from other co-authors, and all authors revised and approved the submitted version.

## Author Information

Correspondence and requests for materials should be addressed to X.L., L.S., or S. T.

## Data availability

The 3D cryo-EM density map of the Naproxen-bound AtPIN1 has been deposited in the Electron Microscopy Data Bank (EMDB, https://www.ebi.ac.uk/emdb/) under the accession number EMD-#####. Coordinates for the Naproxen-bound structure model has been deposited in the Protein Data Bank (PDB, https://www.rcsb.org/) under the accession code ####. Source data for Figures 1, 2, 3, and 4, and Extended Figures 1, 2, 3, 4, and 5 are provided with the paper. All data necessary to evaluate the conclusions in the paper or the supplementary materials are available from the corresponding authors upon request.

## Methods

### Plant materials and growth conditions

*Arabidopsis thaliana* (L.) lines used in this study were in the Columbia-0 (Col-0) ecotype background. The marker lines *DR5rev::GFP* (Friml et al., 2003) and *DR5v2::tdTomato;DR5rev::n3GFP* (Liao et al., 2015) were described previously.

For phenotyping of seedlings or pharmacological experiments, surface-sterilized seeds (using 75% ethanol) were sown on 0.5× Murashige and Skoog (MS) medium supplemented with 1% (w/v) sucrose and 0.8% (w/v) phytoagar (pH 5.9), stratified at 4 °C for 2 d and then grown in a growth chamber at 21 °C with a long-day photoperiod (16 h light-8 h dark). For the dark treatment, the plates were covered with aluminum foil. For the phototropic response assay, etiolated seedlings (aged 4 d) were exposed to light for 6 h, and were then exposed to a directional light source (compact fluorescent lamp, Philips). For microscopic analysis, five-day-old seedlings of different reporter lines were treated with Naproxen and NPA on MS plates.

### Pharmacological treatments

For pharmacological treatments, *Arabidopsis* seeds were sown on MS plates with the indicated chemicals, including Naproxen (Sigma, N8280-5G) and NPA (Sigma, N12507-250MG). After stratification for 2 d at 4 °C, the plates were transferred to a growth chamber under conditions as described in the ‘Plant materials and growth conditions’ section for 5 d, 7 d or 10 d according to different assays. The phenotype was then analyzed using CLSM or ImageJ.

For short-term treatment to study the subcellular localization of *DR5v2::ntdTomato;DR5rev::n3GFP* seedlings (aged 5 d) grown on normal MS plates were transferred to DMSO/Naproxen/NPA-containing MS plates, and turned by 90° for 4 h. Afterwards, the samples were imaged using confocal laser scanning microscopy (CLSM).

### CLSM imaging

Fluorescence imaging was performed using a Zeiss LSM980 CLSM equipped with a GaAsP detector (Zeiss). The manufacturer’s default settings (smart mode) were used for imaging proteins tagged with GFP (excitation, 488 nm; emission, 495–545 nm). All of the images were obtained in 8-bit depth with 2× line averaging. Images were analyzed with the ImageJ software (NIH).

### Statolith starch staining using Lugol solution

Five-day-old *Arabidopsis* Col-0 seedlings treated with Naproxen and NPA (with DMSO as the solvent control) were stained with Lugol solution for 2 min, and were then washed with liquid MS medium for 2 min. The samples were mounted with a clearing solution (30 ml H_2_O, 10 ml glycerol, 80 g chloral hydrate to a total of 100 ml). Finally, root tips were imaged using a DIC microscope (Olympus BX53).

### Imaging and morphological analysis

For physiological experiments, MS plates with *Arabidopsis* seedlings were imaged with a camera (Sony A600 with a macro lens), and then the primary root length, root tip angles, hypocotyl length or growth direction was analyzed with the ImageJ software (NIH). Lateral root numbers were counted directly. Apical hooks of etiolated seedlings were imaged using a stereomicroscope (Nikon, SMZ1500).

### Protein expression and purification

Protein purification process was conducted as described previously (Yang et al., 2022). The DNA sequence of full-length *Arabidopsis thaliana* PIN1 was subcloned into the pCAG vector (Invitrogen) with the N-terminal Flag tag. The HEK 293F cells (Sino Biological Inc.) were cultured in SMM 293T-II medium (Sino Biological Inc.) at 37 °C supplemented with 5% CO_2_. 1.5 mg plasmids and 4 mg inear polyethylenimines (PEI) (Polysciences) were pre-incubated in 50 ml fresh medium for 30 min before adding into one litre cells at density of 3.0 × 10^6^ cells ml^−1^. Then the mixture was added into cell culture followed by 15-min incubation. The transfected cells were collected after 60 h culture and resuspended in the lysis buffer containing 25 mM HEPES pH 7.4, 150 mM NaCl, supplied with 1.3 mg ml^−1^ aprotinin, 5 mg ml^−1^ leupeptin, 0.7 mg ml^−1^ pepstatin, and 1 mM phenylmethylsulfonyl fluoride (PMSF). Then the suspension was incubated in the buffer with additional l.5 % (w/v) n-dodecyl-β-D-maltopyranoside (DDM, Anatrace) at 4 °C for 2 h. After centrifugation at 14,000 rpm for 60 min, the supernatant was incubated with the anti-Flag M2 affinity gel (Sigma) at 4 °C for 45 min. The resin was rinsed 3 times with 10 ml buffer each time containing 25 mM HEPES, pH 7.4, 150 mM NaCl, 0.06% (w/v) glyco-diosgenin (GDN) (Anatrace). The protein was eluted with wash buffer plus 200 µg/ml Flag peptide. To prepare PIN1 and the sybody (sybody-21 used for final structure determination) complex, purified PIN1 and sybody-21 were mixed together with a molar ratio of ∼1:3 and then applied to size-exclusion chromatography using a Superose 6 Increase column (GE Healthcare) in buffer containing 25 mM HEPES pH 7.4, 150 mM NaCl and 0.02% GDN. Peak fractions were pooled and concentrated to approximately 6 mg ml^−1^ for structural studies.

### Sample preparation and cryo-EM data acquisition

For the Naproxen-bound PIN1 cryo-EM sample preparation, 1 mM Naproxen (Aladdin) was first incubated with the protein on ice for 1 hour, then 4 µl of the sample was applied to glow-discharged holey carbon grids (Quantifoil Cu R1.2/1.3, 300 mesh). The grid was blotted for 4s and then plunged into liquid ethane using a Vitrobot Mark IV (FEI) at 8 °C and 100% humidity. The dataset was collected on a 300 kV Titan Krios cryo-EM (Thermo Fisher Scientific) with a nominal magnification of 81,000×. Images were recorded by a K3 direct electron detector positioned after a quantum energy filter (Gatan) in super-resolution mode with a pixel size 0.55 Å by EPU software (Thermo Fisher Scientific). A total of 2414 images were collected with defocus values varying from -1.5 to -2.3 µm. Each image was acquired with an exposure time of 3s and dose-fractionated to 32 frames with total dose of 50 e^−^ Å^−2^.

### Image processing

The data processing workflow is presented in Extended Figure 5g. All micrographs were first imported into Relion 3.1 for image processing (Zivanov et al., 2018). The beam-induced motion correction and dose weighting were performed using MotionCor2 (Zheng et al., 2017). Micrographs were then imported in cryoSPARC for all subsequent image processing tasks (Punjani et al., 2017). Contrast transfer function (CTF) estimations were done with implemented patch CTF. For the structure analysis, 115,7541 articles were automatically picked for two-dimensional (2D) classification. 365,141 particles were selected after 2D classification, and then subjected to ab-initial 3D classification into 5 classes. The beast class was selected for non-uniform refinement and a map at 4.2 Å was obtained. Further ab-initio reconstructions and hetero-refinement were carried out to improve the resolution. The final 3D cryo-EM density map was reconstructed and refined at 3.7 Å, as estimated with the gold-standard FSC at a 0.143 criterion with a high-resolution noise substitution method (Rosenthal and Henderson, 2003; Chen et al., 2013). Local resolution estimation was performed using cryoSPARC.

### Model building

The 3.7 Å reconstruction map for Naproxen-bound PIN1 were used for model building in COOT (Emsley et al., 2010). The previously solved structure of apo PIN1 (PDB ID: 7Y9T) was docked into the cryo-EM map using UCSF Chimera to generate the initial model. The built model was refined with iterative cycles of manual refitting in COOT (Emsley et al., 2010) and real space refined using PHENIX with secondary structure and geometry restraints (Adams et al., 2010). Statistics of the 3D reconstruction and model refinement can be found in Extended Data Table 1.

### Surface plasmon resonance

The SPR analysis was performed using a Biacore 8K system (Cytiva) system at 25 °C with a flow rate of 30 µl min^−1^. Purified wild-type PIN1 proteins were immobilized onto the series S CM5 sensor chips (Cytiva) by amine-coupling chemistry. Naproxen at different concentrations were flowed over the chip surface in the pH 7.0 running buffer, consisting of 25 mM HEPES, pH 7.0, 150 mM NaCl, 0.01% GDN and 5% DMSO. Data were analyzed with the Biacore Insight Evaluation Software Version 3.0.12 using steady state affinity binding model.

### Cell-based auxin efflux assays

Auxin efflux assays followed a general protocol as reported previously (Yang et al., 2022). In brief, PIN1 and *A. thaliana* D6PK (Uniprot accession code: Q9FG74) were subcloned into the pCAG vector, respectively. HEK 293F cells at a density of 1.5×10^6^ cells mL^−1^ were co-transfected with PIN1 or the empty vector and D6PK at a mass ration of 3:1. After 24h transfection, cells were harvested by centrifugation and resuspended in the buffer consisting of PBS citrate buffer, pH 5.5 (10 mM Na_2_HPO_4_, 1.8 mM KH_2_PO_4_, 2.7 mM KCl, 137 mM NaCl, pH adjusted by citric acid anhydrous), 50 µM NPA (Sigma) or Naproxen for 5min, and the DMSO as the control. Then cells were loaded in PBS citrate buffer, pH 5.5, 40 nM [^3^H]IAA (specific activity 25 Ci mmol^−1^, American Radiolabeled Chemicals) and 50 µM inhibitors. After 5 min incubation, cells were washed by centrifugation and resuspended with [^3^H]IAA-free PBS citrate buffer. A 500 µL aliquot of cell suspension containing 3×10^6^ cells were taken immediately after resuspension (defined as the zero time point) and 10min time points. Cells were centrifuged and washed twice with 1 mL ice-cold PBS buffer, pH 7.4, and resuspended with the same buffer plus 1% Triton X-100. Finally, 2ml scintillation fluid was added for scintillation counting. The radioactivity in cell extracts were counted using liquid scintillation counting (Tri-Carb 2910TR, PerkinElmer). [^3^H]IAA retention is presented as residual radioactivity measured at 10 min relative to 0 min. All the experiments were repeated three times independently.

### Quantification and statistics

Most experiments have been repeated at least three times independently, with similar results obtained. To measure hypocotyl length, primary root length, root-tip angles, apical hooks and hypocotyl growth direction, photographs were analyzed using ImageJ (https://imagej.nih.gov/ij/download.html). Fluorescence intensity of CLSM images was quantified by Fiji (https://fiji.sc/). Data visualization and statistics were mostly performed with GraphPad Prism 8. For bending curvatures of roots or hypocotyls, polar bar graphs were generated by Origin 2021.

