## Supplemental Figures and Tables for "Chemical inhibition of PIN auxin transporters by the anti-inflammatory drug Naproxen"

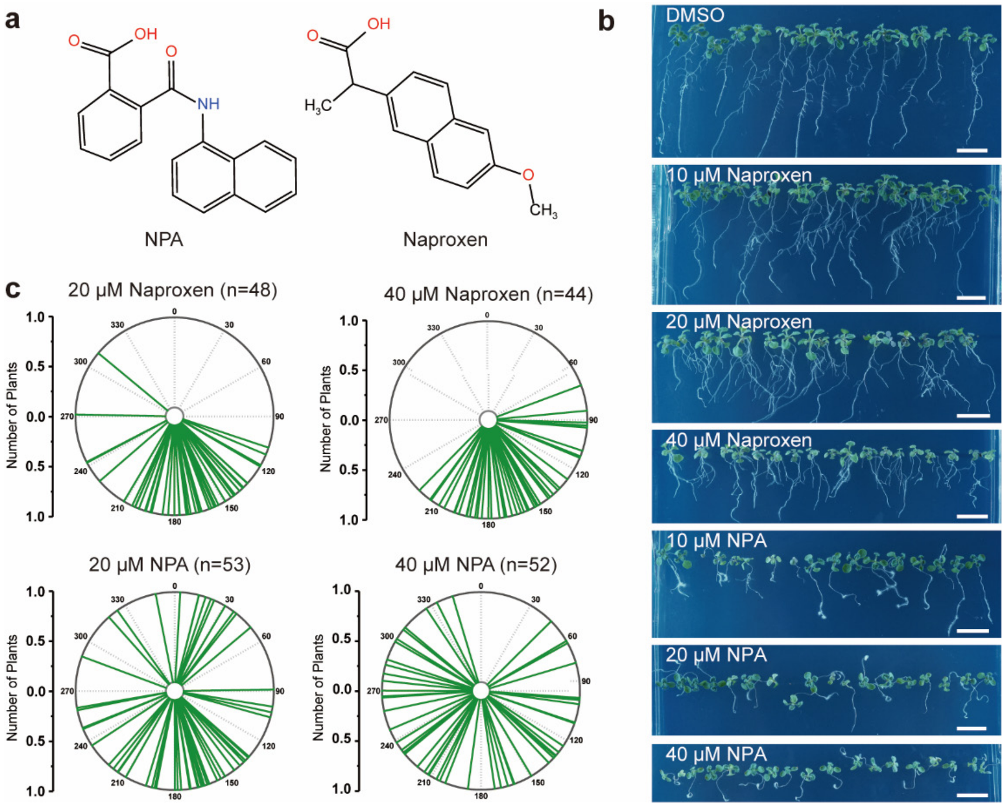


**Extended Data Fig.1. Naproxen modulates *Arabidopsis* root development.**

**a,** Chemical structures of Naproxen and NPA, illustrated using the KingDraw software.

**b**, Representative images showing the morphological changes of 10-day-old Col-0 seedlings grown on MS media supplemented with Naproxen and NPA at indicated concentrations. Scale bars, 1 cm.

**c**, Naproxen and NPA interfered with root gravitropism. Seven-day-old Col-0 seedlings grown constantly on MS medium supplemented with DMSO, Naproxen and NPA, as indicated. Each line represents the root tip angle of 1 individual seedling in polar bar charts.

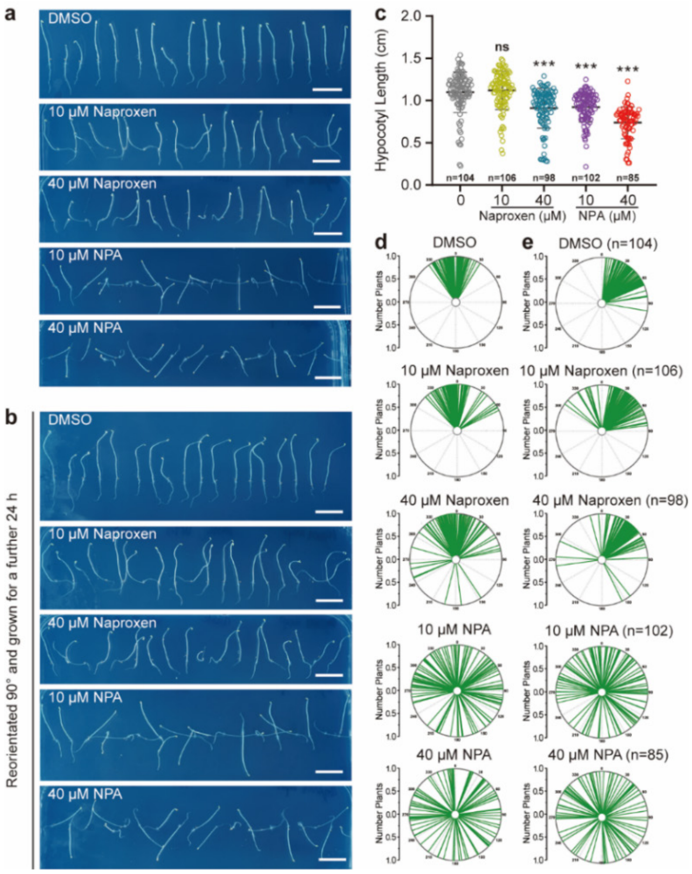


**Extended Data Fig.2. Naproxen impairs gravitropism of *Arabidopsis* seedlings under dark.**

**a**, Representative images showing the morphological changes of etiolated Col-0 seedlings (aged 4 d) grown on MS media supplemented with Naproxen and NPA at indicated concentrations. Scale bars, 1 cm.

**b**, The morphological changes of etiolated Col-0 seedlings (aged 4 d) with Naproxen and NPA were reorientated 90° and grown for a further 24 h. Scale bars, 1 cm.

**c**, The effect of Naproxen and NPA inhibiting hypocotyl elongation under dark conditions. Four-day-old etiolated Col-0 seedlings. DMSO is the solvent control. $p$ values were calculated by comparing different treatments to DMSO with an unpaired $t$ test with Welch's correction, ***$P$<0.001, ns, no significance.

**d** and **e**, Naproxen and NPA interfered with hypocotyl gravitropism. Four-day-old etiolated Col-0 seedlings grown constantly on MS medium supplemented with DMSO, Naproxen and NPA, as indicated. Each line represents the hypocotyl angle of 1 individual seedling in polar bar charts. **e**, Etiolated Col-0 seedlings (aged 4 d) were gravistimulated by 90º reorientation, and hypocotyl inclination was measured at 24 h.

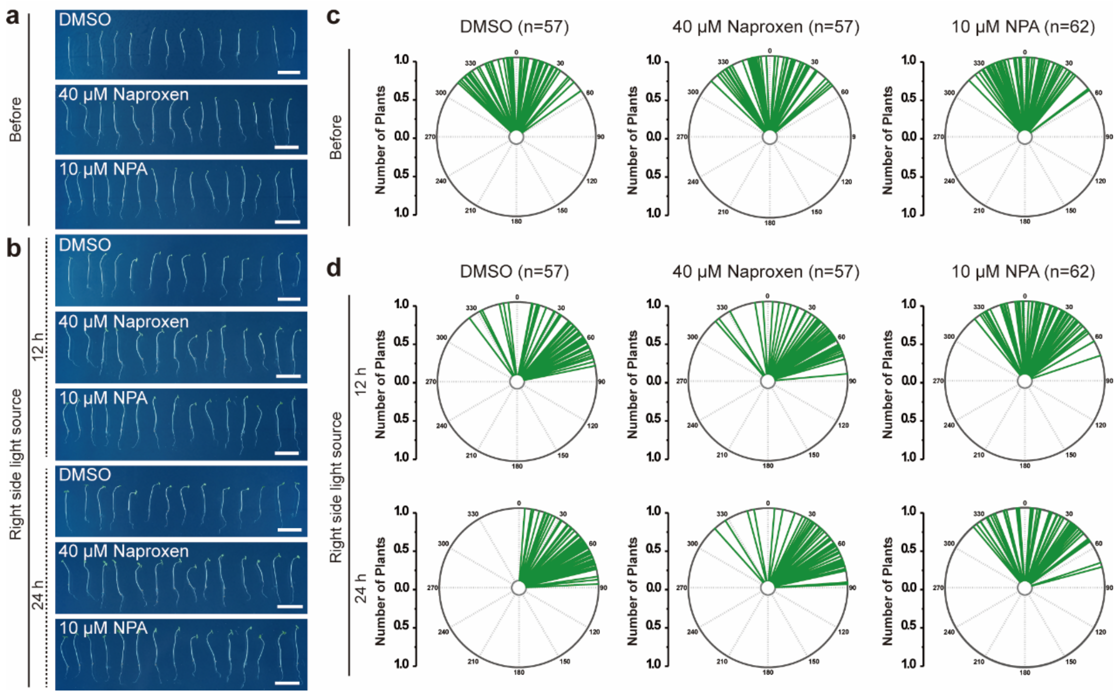


**Extended Data Fig.3. Naproxen treatment compromised the phototropic response of *Arabidopsis* seedlings.**

**a** and **b**, Representative images showing the morphological changes of etiolated Col-0 seedlings (aged 4 d) grown on MS media supplemented with Naproxen and NPA at indicated concentrations. Scale bars, 1 cm. **b**, Etiolated Col-0 seedlings (aged 4 d) were exposed to light for 6 h and further exposed to a directional light source (from the right side) for 12 h and 24 h.

**c** and **d**, The bending angles of the hypocotyls are shown as polar bar charts. Each line represents the hypocotyl angle of 1 individual seedling in polar bar charts.

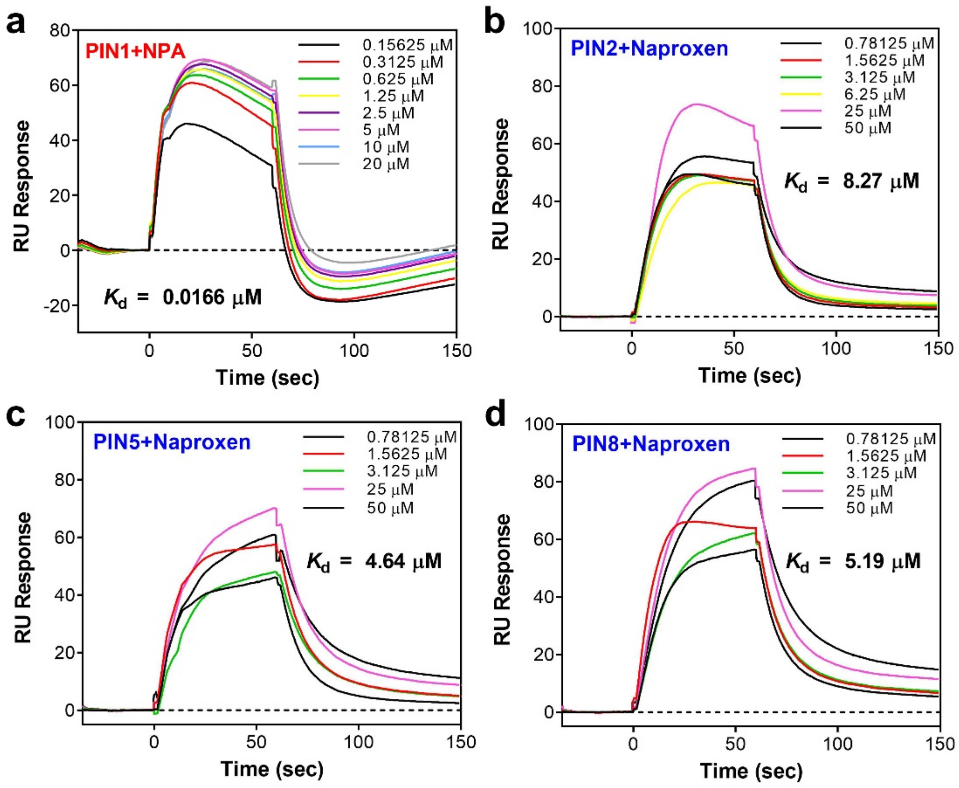


**Extended Data Fig.4. Binding between PIN1/PIN2/PIN5/PIN8 and NPA or Naproxen.**

**a,** The binding affinity between PIN1 and NPA as examined by SPR. The $K_d$ value is 0.0166 uM.

**b-d,** The binding affinity between PIN2, PIN5 or PIN8 and Naproxen as examined by SPR, respectively. The results suggest that Naproxen can bind all PINs examined with similar binding affinities.

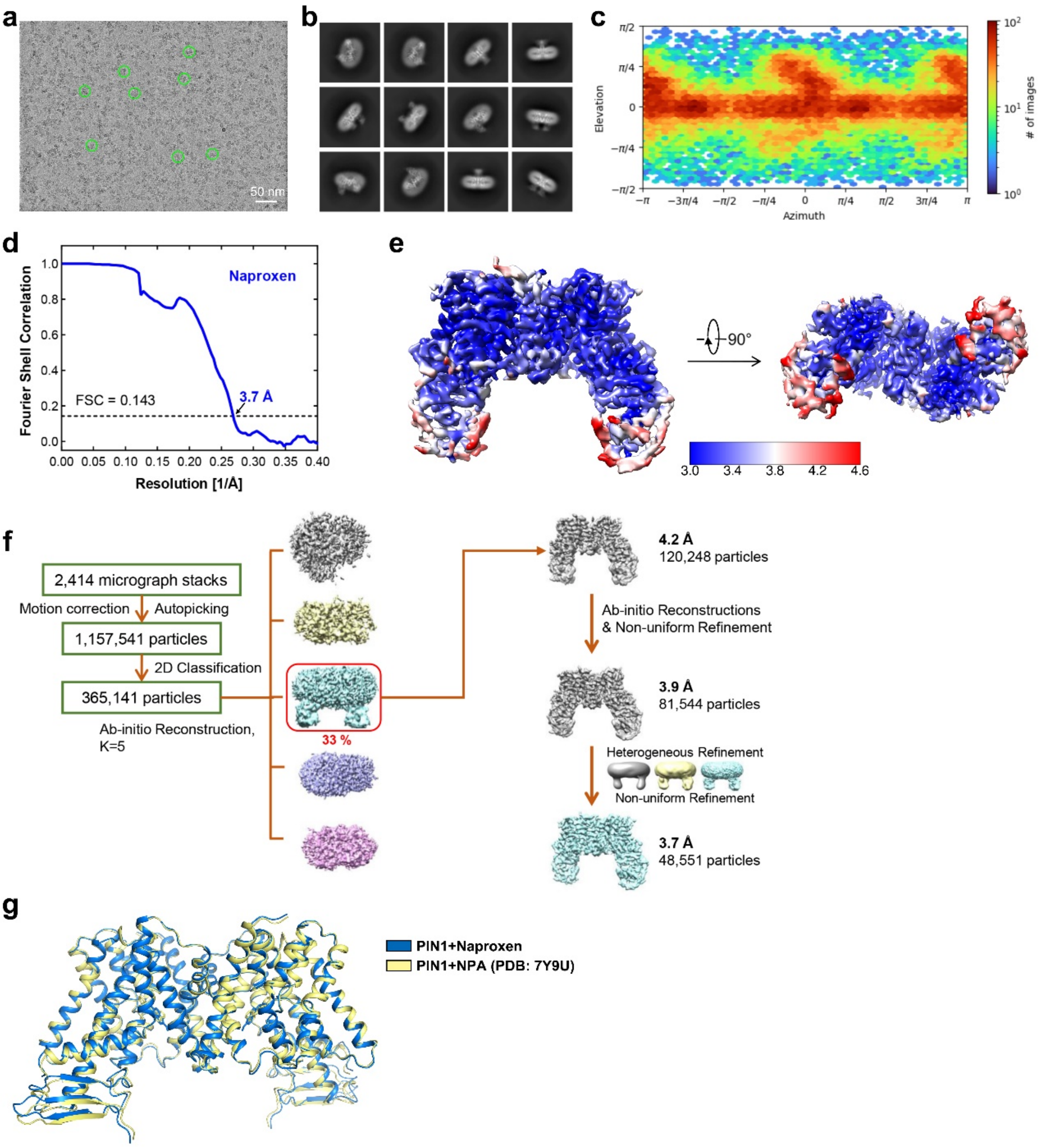


**Extended Data Fig.5. Representative raw micrograph of Naproxen-bound PIN1 and flowchart of the image processing procedure.**

**a,** Representative raw micrograph of Naproxen-bound PIN1 plus sybody-19.

**b,** Representative 2D classes of Naproxen-bound PIN1 plus sybody-19.

**c,** Angular distribution histogram of the final Naproxen-bound PIN1 plus sybody-19 reconstruction.

**d,** The gold-standard Fourier shell correlation curves for the overall map of Naproxen-bound PIN1 plus sybody-19.

**e,** The local resolution estimation of Naproxen-bound PIN1 plus sybody-19.

**f,** Flowchart of the image processing procedure of Naproxen-bound PIN1 structure.

The average resolution for the final reconstruction is estimated to be 3.7 Å.

**g,** Structural alignment of the Naproxen-bound and NPA-bound structures of PIN1. The Naproxen-bound PIN1 structure is shown in marine and the NPA-bound structure (PDB code: 7Y9U) is shown in yellow.

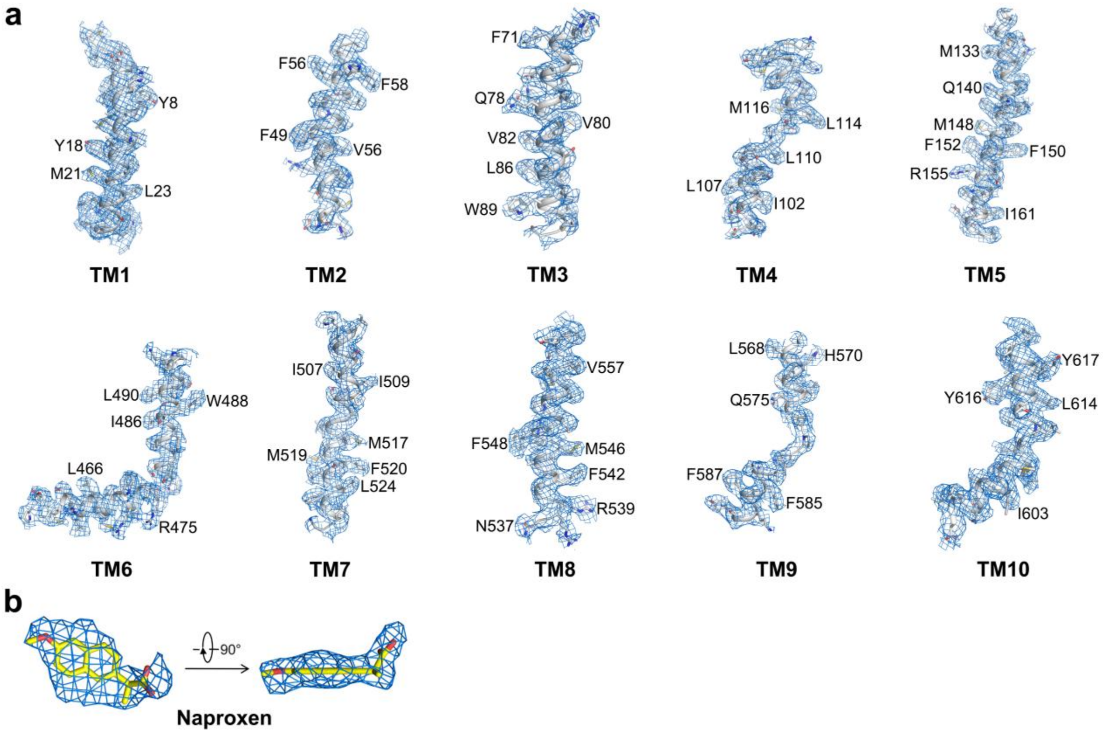


**Extended Data Fig.6. EM density segments of ten transmembrane helices of the Naproxen-bound PIN1 map.**

**a,** EM density segments of ten transmembrane helices of the Naproxen-bound PIN1 map. Residues with notable side chains are labeled aside.

**b,** EM density segments of Naproxen shown in two perpendicular views.

**Extended Data Table 1. Statistics of cryo-EM data collection, processing, model refinement and validation.**

| **Data Collection** | ***At*PIN1+Naproxen** |
|---|---|
| EM equipment | FEI Titan Krios |
| Voltage (kV) | 300 |
| Detector | Gatan BioQuantum K3 |
| Pixel size (Å) | 1.1 |
| Electron dose ($e^-$/Å$^2$) | 50 |
| Defocus range (μm) | -1.5 ~ -2.3 |
| **3D Reconstruction** | |
| Software | RELION, cryoSPARC |
| Number of Particles | 48,551 |
| Symmetry | C2 |
| Map Resolution (Å) | 3.7 |
| FSC Threshold | 0.143 |
| **Model Refinement** | |
| Map Sharpening B-factor (Å$^2$) | -154.7 |
| Model Resolution (Å) | 3.8 |
| FSC Threshold | 0.5 |
| Protein residues | 988 |
| Side chains | 988 |
| Ligands | 2 |
| CC mask | 0.83 |
| **Validation** | |
| R.m.s. Deviations | |
| Bond lengths (Å) | 0.006 |
| Bond angles (°) | 1.205 |
| MolProbity Score | 1.50 |
| All-atom Clashscore | 3.08 |
| Rotamer Outliers (%) | 0.00 |
| Ramachandran plot | |
| Favored (%) | 94.06 |
| Allowed (%) | 5.94 |
| Outliers (%) | 0 |
